# Single-cell immune profiling reveals the impact of antiretroviral therapy on HIV-1-induced immune dysfunction, T cell clonal expansion, and HIV-1 persistence *in vivo*

**DOI:** 10.1101/2021.01.27.428491

**Authors:** Jack A. Collora, Delia Pinto-Santini, Siavash Pasalar, Neal Ravindra, Carmela Ganoza, Javier Lama, Ricardo Alfaro, Jennifer Chiarella, Serena Spudich, David van Dijk, Ann Duerr, Ya-Chi Ho

**Affiliations:** Department of Microbial Pathogenesis, Yale University School of Medicine, New Haven, CT 06519, USA; Vaccine and Infectious Disease & Public Health Science Divisions, Fred Hutchinson Cancer Research Center, Seattle, MA, 98109, USA; Department of Internal Medicine (Cardiology), Yale School of Medicine, New Haven, CT 06520, USA; Department of Computer Science, Yale University, New Haven, CT 06520, USA; Asociación Civil Impacta Salud y Educación, Lima, 15063, Perú; Centro de Investigaciones Tecnológicas Biomédicas y Medioambientales (CITBM), Lima, 07006, Perú; Department of Neurology, Yale University School of Medicine, New Haven, CT 06519, USA

## Abstract

Despite antiretroviral therapy (ART), HIV-1 persists in proliferating T cell clones that increase over time. To understand whether early ART affects HIV-1 persistence *in vivo*, we performed single-cell ECCITE-seq and profiled 89,279 CD4^+^ T cells in paired samples during viremia and after immediate versus delayed ART in six people in the randomized interventional Sabes study. We found that immediate ART partially reverted TNF responses while delayed ART did not. Antigen and TNF responses persisted despite immediate ART and shaped the transcriptional landscape of CD4^+^ T cells, HIV-1 RNA^+^ cells, and T cell clones harboring them (clone_HIV-1_). Some HIV-1 RNA^+^ cells reside in the most clonally expanded cytotoxic T cell populations (GZMB and GZMK Th1 cells). Clone_HIV-1_^+^ were larger in clone size, persisted despite ART, and exhibited transcriptional signatures of antigen, cytotoxic effector, and cytokine responses. Using machine-learning algorithms, we identified markers for HIV-1 RNA^+^ cells and clone_HIV-1_^+^ as potential therapeutic targets. Overall, by combining single-cell immune profiling and T cell expansion dynamics tracking, we identified drivers of HIV-1 persistence *in vivo*.

---

Despite effective antiretroviral therapy (ART), HIV-1-infected CD4^+^ T cells not only persist lifelong^1,2^ but also proliferate *in vivo*^3-7^. More than 50% of the HIV-1 latent reservoir is maintained by clonal expansion of infected cells^3-5^. The proliferation of HIV-1-infected cells is a major barrier to cure. The proliferation of HIV-1-infected cells is driven by antigen stimulation^8-11^, homeostatic proliferation^12^, and HIV-1-driven proliferation gene expression^6,7,13^. Understanding the immune programs promoting the clonal expansion of HIV-1-infected cells is critical for stopping the proliferation of HIV-1-infected cells.

ART suppresses HIV-1 plasma viral load to clinically undetectable levels. Early ART initiated within 6 months of infection, as opposed to delayed ART initiated after 6 months of infection, significantly reduces chronic immune activation^14^. However, early ART may not fully restore immune function to the levels in uninfected individuals^15-17^. ART neither kills infected cells nor inhibits HIV-1 viral protein production from the existing infected cells. Therefore, HIV-1-infected cells can continue to produce viral antigens^18,19^ and induce immune activation and exhaustion^19,20^. We postulate that both antigen stimulation and systemic immune activation promote the proliferation of HIV-1-infected cells. Given that people treated with immediate ART have a shorter duration of HIV-1 viremia, shorter duration of overt HIV-1 antigen exposure, and lower levels of immune activation^14^, we test our hypothesis by comparing the immune programs of CD4^+^ T cells, HIV-1-infected cells, and T cell clonal expansion dynamics in people treated with immediate ART versus delayed ART. In particular, we want to examine the drivers of HIV-1 persistence both for the HIV-1-infected cells and the T cell clones harboring them, given that T cells having the same T cell receptor (TCR) sequence respond to the same antigen stimulation and immune activation. We propose to identify dysregulated immune programs that promote the persistence of HIV-1-infected cells and identify cellular markers of HIV-1 RNA^+^ cells and T cell clones harboring them for therapeutic targeting.

Several challenges prevent mechanistic understanding of HIV-1 persistence in people with HIV-1. First, inducible HIV-1-infected cells in ART-treated individuals are extremely rare, accounting for <0.001% of CD4^+^ T cells in the peripheral blood^21-23^. Second, there are no known cellular markers that exclusively select for heterogenous HIV-1-infected T cells, given the differences^24^ in polarization^25^, activation state^26^, memory differentiation^27-29^, and immune exhaustion states^30^. Third, while HIV-1-infected cells can be isolated using viral markers such as HIV-1 RNA expression^13^ or Env protein expression^31^, these methods require *ex vivo* activation and thus cannot evaluate the *in vivo* status of infected cells. Advancements in single-cell technologies enabled high-dimensional immune profiling to dissect the heterogeneous states of immune cells, high-resolution capture of rare cells, T cell clonality, and identification of upstream drivers of immune dysregulation^32-37^. Further, computational techniques including supervised and unsupervised machine learning, network analysis, and statistical methodologies enable confident identification of higher-fidelity predictors of different cellular states from the sparse and highly complex single-cell multi-modal data^38-43^.

Here, we combined a translational approach using paired and longitudinally archived blood samples from a prospective, randomized, and interventional Sabes study, single-cell multi-omics immune profiling, and machine learning analysis to capture HIV-1-infected cells without *ex vivo* activation. We tracked T cell clonal expansion dynamics over time, identified T cells harboring the rare HIV-1-infected cells, identified immune programs driving HIV-1 persistence, and identified cellular markers for HIV-1 RNA^+^ cells and T cell clones harboring them as therapeutic targets.

## Results

### Single-cell multi-omics of CD4^+^ T cells from the randomized and interventional Sabes study identifies the impact of immediate versus delayed ART on HIV-1-induced immune dysfunction

To understand why people treated with immediate versus delayed ART have different immune dysfunction^15-17^, we compared the immune profiles and T cell clonal expansion dynamics of CD4^+^ T cells from people with acute HIV-1 infection treated with immediate versus delayed ART. We used longitudinally archived blood samples from the Sabes Study in which HIV-1 infection was prospectively tested monthly for early detection and treatment stratification^44,45^. Using detailed histories of HIV-1 testing, including date, type and result of test, we calculated the estimated date of detectable infection (EDDI) using an established algorithm^46^. We examined the single-cell transcriptional landscape of six people with acute HIV-1 infection treated either with immediate ART (three individuals receiving ART 22 – 47 days after EDDI) versus delayed ART (three individuals receiving ART 187 – 207 days after EDDI) (Fig. 1a, Supplementary Fig. 1, Supplementary Table 1). Paired blood samples during viremia (20 – 48 days after EDDI) and after one year of ART (172 – 384 days of suppression) from each of the six individuals were profiled to examine the cellular transcriptional landscape. Two biological sex-matched uninfected individuals were recruited for comparison.

**Fig. 1.**
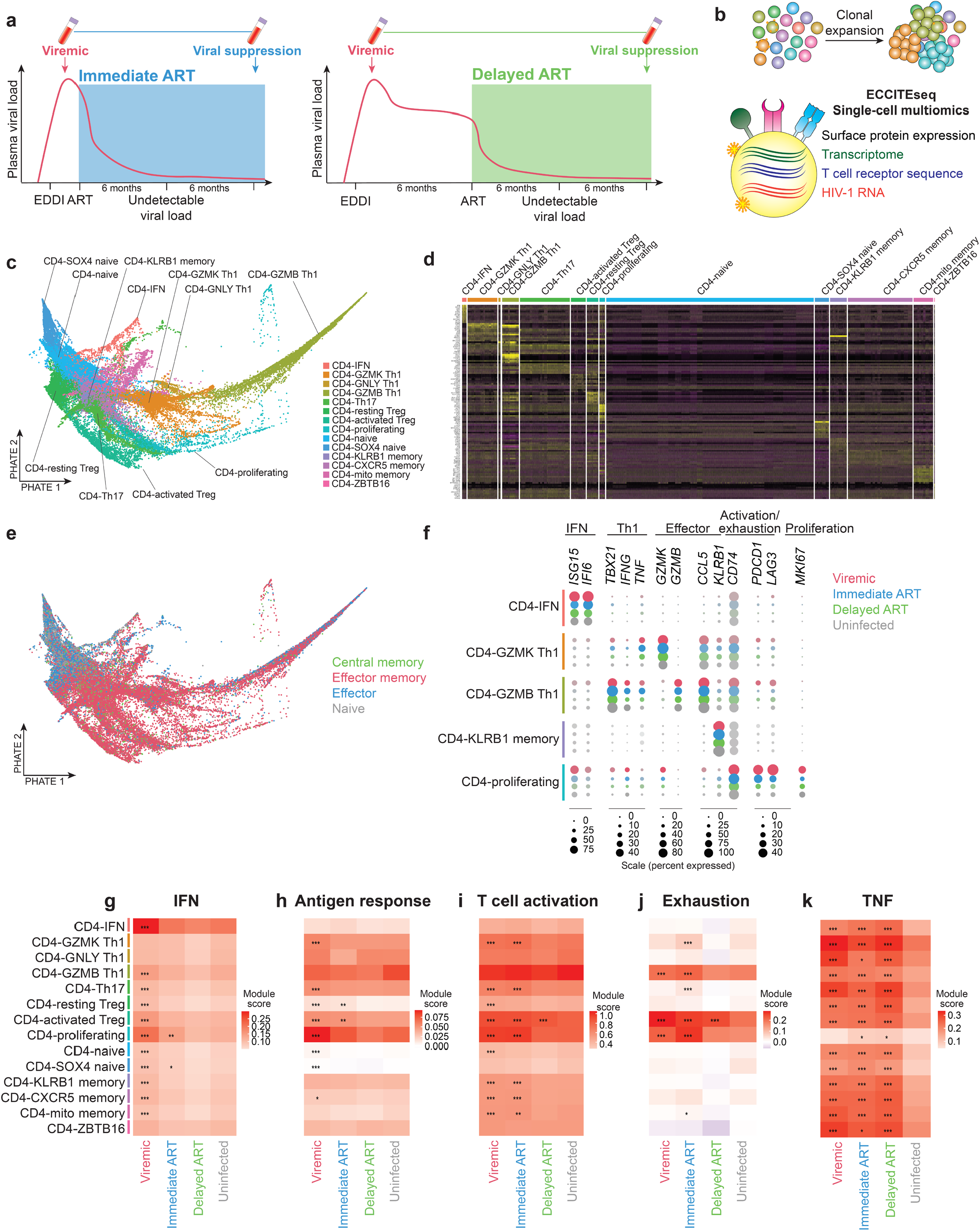
Single-cell immune profiling of CD4^+^ T cells of paired longitudinal blood samples during viremia and after immediate versus delayed initiation of suppressive ART. **a**, Study design. Participants enrolled the Sabes study were prospectively tested monthly for HIV-1 infection using antibody detection, antigen detection, and viral RNA quantification. Paired CD4^+^ T cells during acute viremia and after suppressive ART from six people with HIV-1 enrolled in Sabes and subsequently followed in the Merlin study were used for immune profiling. Among them, three individuals received immediate ART and three individuals received delayed ART (24 weeks after EDDI). CD4^+^ T cells from two sex-matched uninfected individuals were used as controls. **b**, Single-cell ECCITE-seq captures surface protein expression, cellular transcriptome (n = 52,473, 13,569, 23,237, and 33,406 cells in the viremic, immediate ART, delayed ART, and uninfected conditions respectively), HIV-1 RNA, and TCR sequence within the same single cell, allowing clonal expansion dynamics tracking in longitudinally archived samples. **c**, PHATE plot of all 122,685 CD4^+^ T cells from HIV-1-infected and uninfected individuals depicts 14 clusters of CD4^+^ T cells defined by transcriptome. Clusters were labeled with inferred cell type based on transcriptome signatures and surface protein expression. **d**, Heatmap of top 10 differentially expressed genes in each cluster. Heat map marker genes were identified by Wilcoxon rank-sum test with a cutoff of 0.25 average log fold change and an adjusted *P* value cutoff of 0.05. **e**, PHATE plot of memory phenotypes defined by surface CD45RA and CCR7 expression. CD45RA and CCR7 positivity was determined by >90^th^ percentile of the expression level of isotype barcoded surface protein staining antibody controls. **f**, The expression profile of genes involved in type I IFN responses, Th1 polarization, cytotoxic effector functions, T cell activation, and exhaustion phenotypes of CD4^+^ T cell clusters of interest. Color intensity indicates level of expression. Dot size indicates the percentage of cells expressing transcript. **g–k**, Mean module scores of predefined gene sets involved in different immune responses during viremia, viral suppression after immediate ART, viral suppression after delayed ART, and uninfected conditions. **g**, IFN response module was defined by top 30 genes correlating with *ISG15*. **h**, Antigen response module was defined by Goldrath antigen response^52^. **i**, T cell activation module was defined by top 30 genes correlating with *CD74* (HLA-DR antigens-associated invariant chain) expression. **j**, Exhaustion module was defined by top 30 genes correlated with *PDCD1* (PD-1) expression. **k**, TNF response module was defined by Phong TNF response^53^. *P* values were determined by Wilcoxon rank-sum test comparing HIV-1 infected conditions and uninfected conditions. *, *P* <0.01; **, *P* <0.001; ***, *P* <0.0001.

To characterize the heterogeneous functional states of CD4^+^ T cells, to track T cell clonal expansion dynamics, and to identify the rare HIV-1-infected cells, we used Expanded CRISPR compatible Cellular Indexing of Transcriptomes and Epitopes by Sequencing (ECCITE-seq^47^) to profile the cellular transcriptome, surface protein markers, TCR sequence, and HIV-1 RNA within the same single cells (Fig.1b). A total of 52,473 CD4^+^ T cells during viremia, 36,806 CD4^+^ T cells during suppressive ART, and 33,406 CD4^+^ T cells from uninfected individuals were examined by single-cell ECCITE-seq (Supplementary Table 2). Batch effects were removed using Seurat^42^ integration, as shown in dimensionality reduction plots by Potential of Heat-diffusion for Affinity-based Trajectory Embedding (PHATE)^48^(Supplementary Fig. 2). The single-cell transcriptional landscape of all 122,685 cells [having a median of 4,144 unique molecular identifiers (UMI) and 1,345 genes per cell] were visualized using PHATE^48^ (Fig. 1c)(Supplementary Table 3).

We identified 14 transcriptome defined clusters. The inferred CD4^+^ T cell phenotypes was determined based on transcriptional markers and surface protein expression (Fig. 1c, 1d, Supplementary Fig. 3 for gene expression violin plots for each cluster, Supplementary Table 4 for genes and proteins that infer the phenotype of each cluster, and Supplementary Table 5 for a list of differentially expressed genes). These included as one interferon (IFN) response cluster, three Th1 clusters, one Th17 cluster, two Treg clusters, one proliferating cell cluster, two naïve cell clusters, one *KLRB1* (CD161) cluster, and one CXCR5 (homing) cluster. Using surface CD45RA and CCR7 protein expression (determined by >90% expression levels of isotype ECCITE-seq antibody controls), we examined the T cell memory phenotype (naïve, central memory, effector memory, and effector)^49^ of each single cell (Fig. 1e). These clusters showed distinct CD4^+^ T cell effector profiles (Fig. 1f, Supplementary Fig. 4). Importantly, in addition to classic Th1 cells expressing high levels of *TBX21* (Tbet), *IFNG, TNF* (tumor necrosis factor), *GZMB* (granzyme B), and *CCL5* (RANTES), we found that the largest population of Th1 cells was the GZMK Th1 cell subset expressing high levels of *GZMK* (granzyme K), *TNF*, and *CCL5* but not *GZMB*. The cytotoxic CD4^+^ T cells expressing granzyme K, previously under-appreciated in HIV-1-infection, have been shown to be important anti-tumor effectors in bladder cancer^50^ and in SARS-CoV-2 infection^51^. In order to identify the underlying phenotype of proliferating cells that can be masked by cell cycle-related gene signatures, we performed sub-clustering of the proliferating cell cluster. Sub-clustering of proliferating cells revealed a subset of cells in viremic and suppressed conditions with increased *GZMK* expression which was absent in uninfected conditions (Supplementary Fig. 5). This indicates that a subset of GZMK expressing cells expressed proliferation and cell-cycle related genes and may have high proliferation potential.

### Immediate ART limits TNF responses but not ongoing antigen response in CD4^+^ T cells

To identify differences of HIV-1-induced immune dysregulation during viremia, and suppression after immediate vs delayed treatment initiation, we examined T cell functional states in each T cell subset by comparing the gene expression profile with pre-defined gene modules, or group of functionally related and co-regulated genes (Fig. 1g – 1k). Using gene set module scoring^54^, we found that IFN responses, particularly in CD4-IFN cells, were upregulated during viremia and returned to baseline after viral suppression (Fig. 1g). Antigen responses and T cell responses peaked during viremia, reflecting overt antigen exposure and T cell activation during acute infection (Fig. 1h, 1i). However, immune exhaustion persisted in some subsets of cells (Fig. 1j), while TNF responses (Fig. 1k) continued in all subsets of cells despite immediate ART or delayed ART.

In addition to using pre-defined gene sets of immune regulatory pathways, we wanted to identify co-regulated genes that can distinguish the immune responses between viremia, immediate ART, delayed ART, and uninfected conditions using *de novo* gene set identification. Using weighted correlation network analysis (WGCNA), we identified 39 modules across individuals which were merged into eight consensus modules based on gene overlap (Supplementary Fig. 6). From eight consensus gene modules we found three modules which differentiated between different conditions across all participants: IFN responses (Fig. 2a–2d), antigen responses (Fig. 2e–2h), and TNF responses (Fig. 2i–2l)(see Supplementary Table 6 for the gene features of each module). The module names of these *de novo* identified gene sets were determined by gene ontology or pathway enrichment identified by Enrichr^55,56^. We compared gene-gene correlation levels between different conditions (Fig. 2a, 2e, 2f, and Supplementary Fig. 7) and examined the module scores on PHATE (Supplementary Fig. 8) and in CD4^+^ T cell clusters that were most affected (Fig. 2b, 2f, 2i, Supplementary Fig. 9). In these gene modules, we identified the enriched pathways of these co-regulated genes (Fig. 2c, 2g, 2k) and upstream regulators of these genes (Fig. 2d, 2h, 2l) using Ingenuity Pathway Analysis (IPA)^57^.

**Fig. 2.**
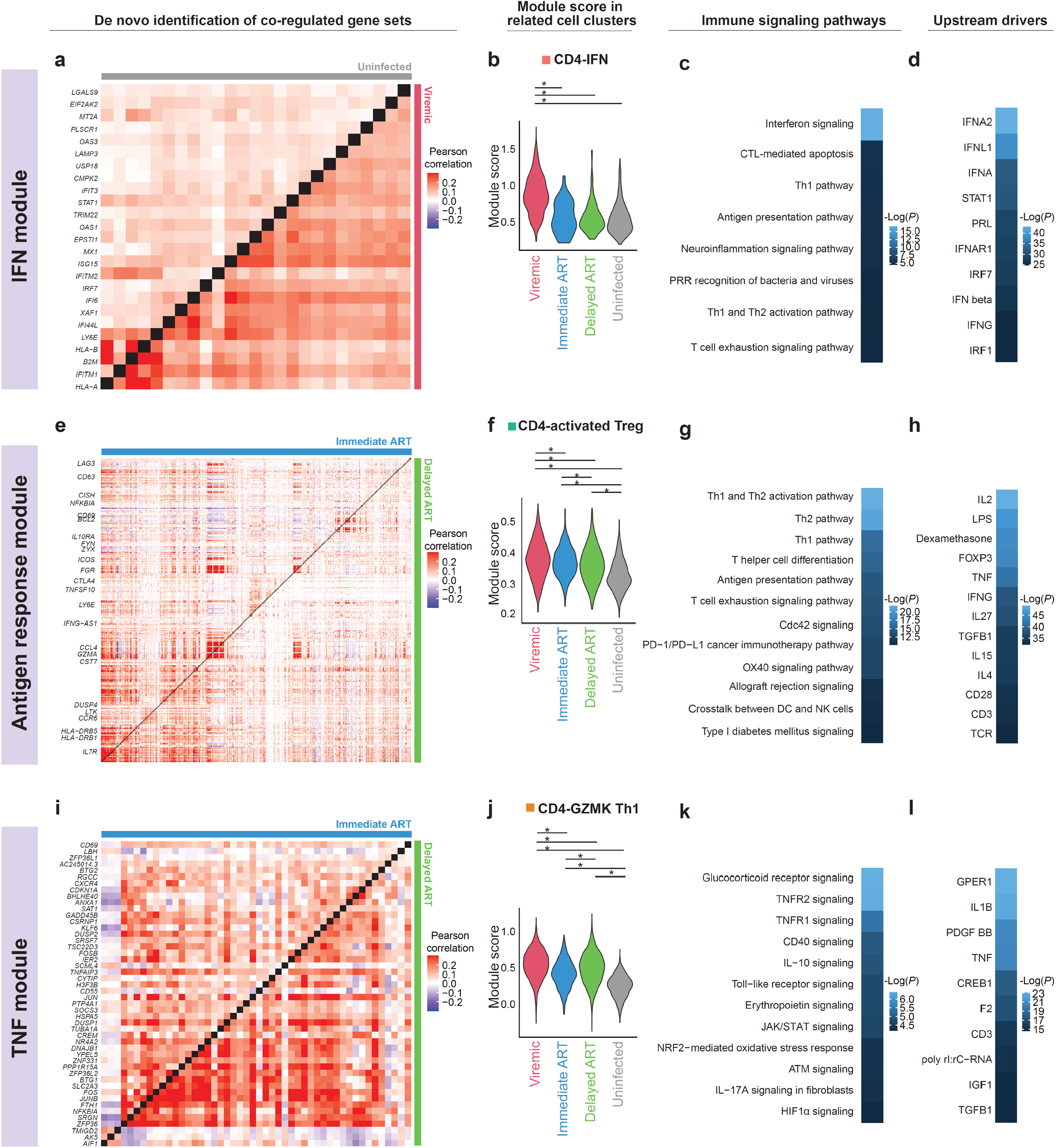
*De novo* identification of co-regulated gene signatures of CD4^+^ T cells during viremia, after immediate ART suppression, after delayed ART suppression, and in uninfected conditions. Weighted correlation network analysis (WGCNA) on all T cells identified a total of three consensus gene modules across all participants that can differentiate between the four conditions. These three gene modules identified *de novo* were IFN responses (a – d, 25 genes), antigen responses (e – h, 348 genes), and TNF responses (i – l, 48 genes). **a, e, i**, Heatmaps of Pearson correlation coefficients of gene expression levels between different conditions. **b, f, j**, Violin plots of module score in clusters showing most prominent differences, such as IFN module score in CD4-IFN cells (n = 1,133 cells), antigen response module score in CD4-activated Tregs (n = 3,073 cells), and TNF module score in GZMK Th1 cells (n = 7,973). *P* values were determined by Wilcoxon rank-sum test. **c, g, k**, Immune pathways significantly enriched in respective modules. **d, h, l**, Upstream regulators driving the transcriptional landscape in respective modules. Immune pathways and upstream regulators were identified using IPA. *P* values were calculated by Fisher’s exact test. *, *P* <0.05.

The IFN response module was characterized by 25 IFN-stimulated genes (ISGs) such as *ISG15, IFI6, IFI44L, IRF7, MX1, IFITM2, IFITM3, STAT1, OAS1, OAS3*, and *XAF1* (Fig. 2a). Consistent with analysis using pre-defined IFN gene sets (Fig. 1g), the upregulation of IFN response module (Fig. 2a) returned to baseline after one year of suppressive ART (Fig. 2b, Supplementary Fig. 8a, Supplementary Fig. 9a). These IFN-stimulated genes were upregulated in acute HIV-1 viremia (Fig. 2b) but also in hyperacute HIV-1 viremia^34^, SIV infection^58^, SARS-CoV-2 infection^59^, and influenza infection^60^, reflecting a shared type I IFN response to acute viral infections. The pathways and upstream regulators are IFN signaling and type I IFNs (Fig. 2c and 2d). Overall, type I IFN responses were induced upon acute HIV-1 infection and returned to baseline after suppressive ART.

The antigen response module included genes involved in T cell exhaustion (*TOX, TOX2, PRDM1* (encoding BLIMP-1), *LAG3*, and *TIGIT*), MHC class II (*CD74* (encoding MHCII invariant chain), *HLA-DRB1*, and *HLA-DRB5*), *ICOS*, and cytotoxic effector molecules (*GZMA, GZMH*, and *PFN1*)(Fig. 2e). This module of 439 genes provides a broader scope of antigen responses and T cell activation than the predefined gene sets (Fig. 1h, 1i). We found that immediate ART slightly reduced antigen responses and T cell activation during HIV-1 infection compared with delayed ART but did not restore antigen responses and T cell activation to levels in uninfected individuals (Fig. 2f, Supplementary Fig. 8b, Supplementary Fig. 9b). The immune pathways involved in this module include Th1 and Th2 activation, T cell differentiation, antigen presentation, and immune exhaustion (Fig. 2g). The upstream regulators driving this persistent T cell activation involved cytokine dysregulation [interleukin-2 (IL-2), TNF, IFNγ, IL-27, tumor growth factor-β (TGFβ), IL-15], lipopolysaccharide (LPS), and TCR stimulation.

The TNF response module included *DUSP1, DUSP2, JUN, JUNB, FOS, FOSB, NFKBIA, ZFP36*, and *BHLHE40* (Fig. 2i). These TNF response genes were upregulated during viremia and persisted during delayed viral suppression (Fig. 2j, Supplementary Fig. 3c). We found that immediate ART significantly reduced TNF responses relative to delayed ART, but not to levels in uninfected individuals (Fig. 2i, 2j, Supplementary Fig. 8c, Supplementary Fig. 9c). The immune pathways involved in this module included glucocorticoid receptor signaling, TNF receptor signaling, OX40 signaling, IL-10 signaling, and JAK/STAT signaling (Fig. 2k). The upstream cytokine regulators driving persistent TNF responses included IL-1β, TNF, and TGFβ (Fig. 2j). Of note, some TNF-response genes (*DUSP1, FOS, FOSB, JUN, JUNB, NFKBIA*) were reported to be upregulated in coronavirus disease 2019 (COVID-19) patients^61^, and in severe COVID-19 but not in severe influenza infection^60^. Further upregulation of TNF/IL-1β-driven inflammatory response was found in hyperacute HIV-1 infection^34^ and in chronically infected untreated HIV-1 infected individuals^62^. Overall, we found that immediate ART slightly reduced antigen responses and T cell activation and significantly reduced TNF responses compared to delayed ART, although this effect is partial. People with HIV-1, despite immediate or delayed ART, may continue to have TNF responses and chronic inflammation.

### HIV-1 RNA^+^ cells have diverse T cell phenotypes

To determine the transcriptional landscape of HIV-1-infected cells, we mapped the transcriptome to autologous HIV-1 genomes and identified 90 HIV-1 RNA^+^ cells, including 81 from viremic samples and 9 from ART-suppressed samples (Fig. 3a). The low number of HIV-1 RNA^+^ cells identified during viremia (0.15%) and after suppressive ART (0.02%) was consistent with the low frequency of HIV-1-infected cells. In these HIV-1 RNA^+^ cells, there were comparable levels of HIV-1 RNA transcripts (normalized to cellular transcripts) in each single cell in viremic samples versus ART-suppressed samples (Supplementary Fig. 10), suggesting that HIV-1-infected cells expressed detectable levels of HIV-1 RNA *in vivo* despite suppressive ART^63^. Of note, cells that are negative for HIV-1 RNA are not necessarily uninfected cells – HIV-1 infected cells in latency or cells harboring transcription-defective HIV-1 may not express HIV-1 RNA. Given the low frequency of total HIV-1 DNA levels in CD4^+^ T cells (<0.1%)^22,23^, >99% of cells that do not have detectable levels of HIV-1 RNA are presumably uninfected. Although HIV-1 RNA positivity does not equal replication competence, these are induced HIV-1-infected cells that are actively producing viral products *in vivo*. There were zero reads mapped to HIV-1 genomes in uninfected individuals. We first examined the normalized frequency of HIV-1 RNA^+^ cells in respect to the size of the respective cluster. We found that a median of 0.42% of GZMK Th1 cells harbored HIV-1 RNA^+^ cells, followed by KRLB1 cells (0.40%), GZMB Th1 cells (0.25%), Th17 cells (0.22%), mitochondria-high cells (0.21%), activated Tregs (0.15%), CXCR5 cells (0.06%), and naïve cells (0.03%)(Fig. 3b). We next examined the number of HIV-1 RNA^+^ cells in each cluster regardless of the size of the cluster. We found that HIV-1 RNA^+^ cells resided in GZMK Th1 cells (14/90 cells, 15.6%), Th17 cells (14/90 cells, 15.6%), CXCR5 memory cells (11/90 cells, 12.2%), naïve cells (10/90 cells, 11.1%), and KLRB1 memory cells (10/90 cells, 11.1%)(see also Supplementary Fig. 11). As previously reported, *KLRB1* (CD161) cells^64^, Th1 cells^25^, Th17 cells^65^, CXCR5^+^ cells^65,66^, and naïve cells^29^ may harbor HIV-1-infected cells, although we found that naïve cells were significantly dis-enriched for HIV-1 RNA^+^ cells (Supplementary Fig. 11a). Overall, we identified GZMK Th1 cells as a previously unappreciated population that harbors HIV-1 RNA^+^ cells.

**Fig. 3.**
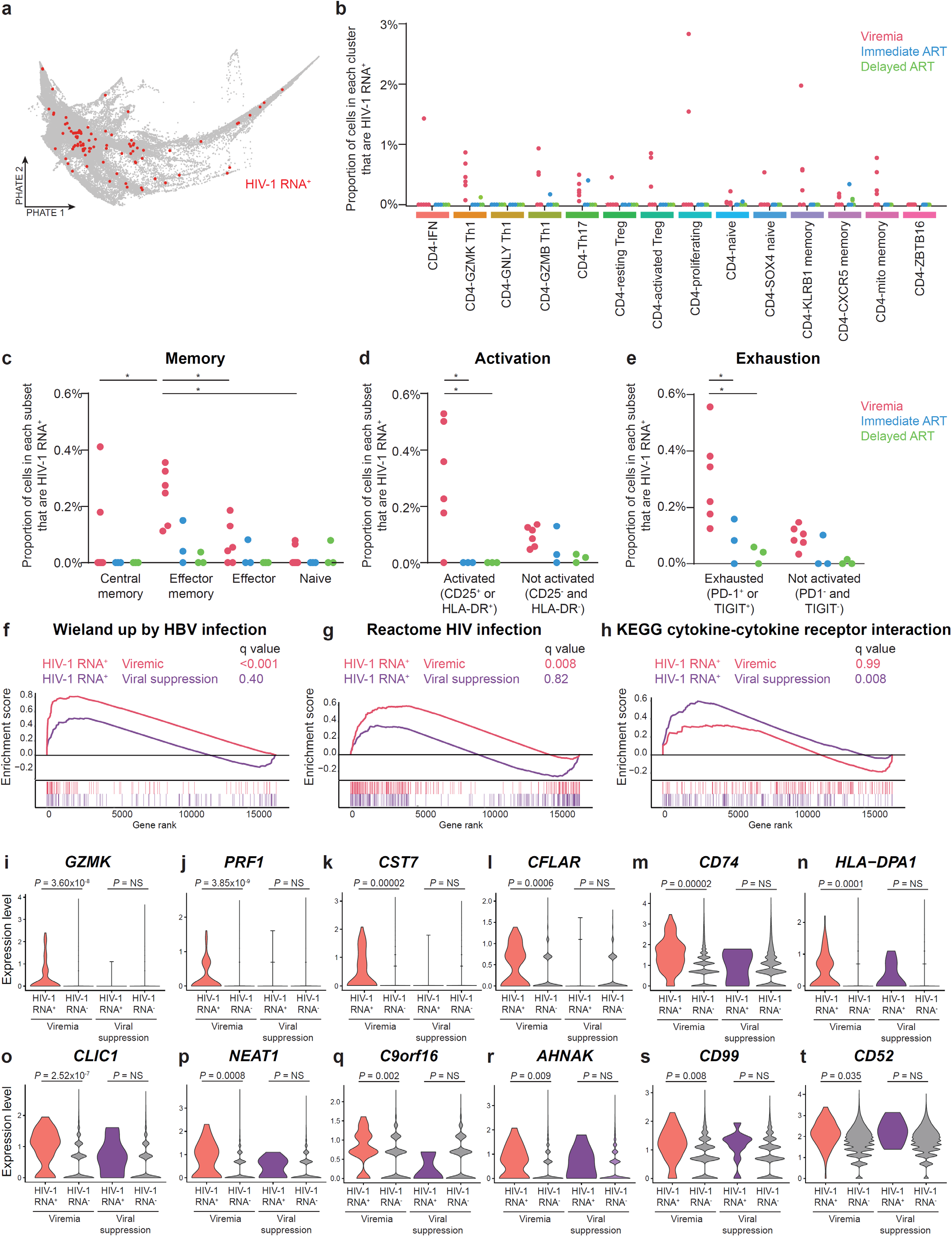
The transcriptional landscape of HIV-1 RNA^+^ cells. **a**, PHATE plot with 90 HIV-1 RNA^+^ cells (including 81 from the viremic condition and 9 from suppressed conditions) labeled in red. **b – e**, Proportion of cells that are HIV-1 RNA^+^ in transcriptome-defined clusters (**b**), memory cell differentiation states (**c**), T cell activation states (**d**), and T cell exhaustion states (**e**). *P* values were determined by two-way ANOVA followed by Tukey’s multiple comparisons test. Surface protein positive staining was determined by >90^th^ percentile of the expression level of isotype barcoded surface protein staining antibody controls. *P* values were determined by two-way ANOVA followed by Tukey’s multiple comparisons test. **f – h**, Identification of immune pathway enrichment in of HIV-1 RNA^+^ cells versus HIV-1 RNA^−^ cells during viremia (**f, g**) and after viral suppression (**h**) using gene set enrichment analysis (GSEA). **i – t**, Markers of HIV-1 RNA^+^ cells identified by bootstrapping. The transcriptome of HIV-1 RNA^+^ cells was compared with HIV-1 RNA^−^ cells (n = 52,392 and 36,797, for viremic and viral suppression respectively) in cluster and individual-matched pools repeatedly for 10,000 times. Genes were selected based on being positive in >95% of test. *P* values were determined by Wilcoxon rank-sum test for 22 genes and corrected by Benjamini-Hochberg procedure. Y axis shows Ln expression levels. *, *P* <0.05.

### HIV-1 RNA^+^ cells are enriched in exhausted and effector memory cells during viremia

Using DNA-barcoded surface protein staining in ECCITE-seq, we examined the memory T cell subsets that were enriched with HIV-1 RNA^+^ cells. We found that the majority of HIV-1 RNA^+^ cells were effector memory cells (CD45RA^−^ CCR7^−^)^28^ during viremia (67/81 cells, 82.7%) and under suppressive ART (7/9 cells, 77.8%), although central memory cells, effector cells, and naïve cells also harbored HIV-1 RNA^+^ cells. The majority of HIV-1 RNA^+^ cells express activation markers during viremia (49/81 cells, 60.5%), but not under suppressive ART (0/9 cells, 0%). We found that around half of HIV-1 RNA^+^ cells (48/81 cells, 59.3%) expressed exhaustion markers (either PD-1 or TIGIT) during viremia and during suppressive ART (4/9, 44.4%). We next examined the proportion of HIV-1 RNA^+^ cells in each subset. We found that effector memory cells, activated cells (expressing CD25 or HLA-DR), and exhausted cells (expressing PD-1 or TIGIT) harbored more HIV-1 RNA^+^ cells during viremia (Fig. 3c– 3e). When we compared the proportion of HIV-1 RNA^+^ cells versus HIV-1 RNA^-^ cells across T cell subsets, we found that HIV-1 RNA^+^ cells were enriched in effector memory cells, dis-enriched in naïve cells during viremia (*P* <0.05)(Supplementary Fig. 11b), and enriched in exhausted T cells (*P* <0.05)(Supplementary Fig. 11d). There is a trend of higher frequency of HIV-1 RNA^+^ cells in activated cells but it did not reach statistical significance (Supplementary Fig. 11c). Consistent with previous studies, we showed that effector memory cell markers^28^, T cell activation markers^26^, and immune exhaustion markers^30,31^ may enrich for a subset of HIV-1-infected cells during viremia.

### TNF and homeostasis cytokine responses shape the transcriptional landscape of HIV-1 RNA^+^ cells during viral suppression

We next examined the gene expression profile of HIV-1 RNA^+^ cells relative to HIV-1 RNA^-^ cells at the same time point both during viremia and after viral suppression. We ranked genes using expression fold change and conducted gene set enrichment analysis (GSEA)^67^ with predefined gene sets^68^. HIV-1 RNA^+^ cells upregulated viral infection-related genes previously identified in the context of HIV-1 infection and hepatitis B virus infection, including *IFI16, IFI27, STAT1, GZMK, GZMA, CCL5, CCR5, CD28, PSMB8*, and *PSMB9* (Fig. 3f, 3g). After ART suppression, the transcriptome of HIV-1 RNA^+^ cells were dominated by cytokine and cytokine receptor interactions, such as cytokines (*TNF, TGFB1,TNFSF13B, TNFSF14, IL7*, and *IL15)* and cytokine receptors (*IL10RA, IL21R, IL2RB, IL15RA, IL21R, IFNAR2*, and *IFNGR1*) (Fig. 3h), suggesting that cytokine dysregulation (such as TNF and TGFβ, Fig. 2k, 2l) and homeostasis common γ chain cytokines (such as IL7 and IL15) during suppressive ART may shape the transcriptome of HIV-1 RNA^+^ cells.

To identify cellular markers that consistently distinguish HIV-1 RNA^+^ cells from HIV-1 RNA^−^ cells from the same individual, we employed a statistical bootstrapping approach to compare HIV-1 RNA^+^ cells to cluster and individual matched pools of HIV-1 RNA^−^ cells repeatedly for 10,000 times. We identified a total of 22 markers (Fig 3i–t, Supplementary Fig. 12, Supplementary Table 7), including molecules related to cytotoxic effectors *GZMK, PRF1* (perforin), and *CST7* (Fig 3i-k). Further, we identified *CFLAR* (c-FLIP)(Fig 3l), a caspase-8 inhibitor that provides resistance to apoptosis, that can be upregulated by Tat expression^69^. We also identified MHC II molecules (*CD74, HLA-DRB1, HLA-DRB5*, and *HLA-DPA1*) (Fig 3m–n, Supplemental Fig. 12 b–d), *CLIC1* (Fig. 3o)(NCC27, a chloride channel involved in cell cycle regulation^70^), *NEAT1* (Fig. 3p)(a long noncoding RNA that restricts Rev-dependent HIV-1 RNA export^71^), and *C9orf16* (Fig. 3q)(of unknown function). Finally, we identified several latency-related genes including *AHNAK* (Fig. 3r, a histone methyltransferase^72^), *CD99* (Fig 3s, upregulated by Tat^73^), and *CD52* (Fig. 3t, a marker of HIV-1 infection during ART^74^).

### The upstream regulators of T cell clonal expansion are antigen stimulation and cytokine dysregulation

TCR sequences can act as barcodes for CD4^+^ T cell clones denoting antigen specificity, due to the sequence diversity generated by genomic rearrangement of the T cell receptor locus. T cells having the same TCR sequence likely originate from the same T cell clone and respond to the same antigen presentation. By examining the T cell repertoire at both viremic and viral suppression time points, we identified cells that were captured as clones (TCR sequence captured two or more times in the same individual) at either time point (Fig. 4a) and tracked the same clones over time. We identified TCR sequences at both the bulk and the single-cell level in separate pools. Bulk TCR sequencing enabled us to achieve a greater depth of TCR capture than single-cell alone and increased our ability to identify clonal T cells and to determine a more accurate clonal frequency. When we examined the transcriptome-defined T cell clusters, we found that GZMB Th1 cells and GZMK Th1 cells were significantly more clonal than other clusters (Fig. 4b, Supplementary Fig. 13). When we examined the memory T cell subsets, we found effector memory cells have the largest proportion of clonal cells (Fig. 4c). To identify cellular transcriptional programs that correlate with T cell clone size, we identified genes having expression levels that positively correlated with T cell clone size by Pearson correlation. Using GSEA, we found that these clone-size correlated genes were strongly involved in antigen responses such as cytotoxic effector expression (*GZMA, GZMB, GZMK, PRF1*), *CCL5, KLRG1, CD74*, and *BHLHE40* in viremic, immediate ART, delayed ART, and uninfected conditions (Fig. 4d). In order to identify regulators of T cell clone size, we used elastic net regression^75^ to learn the relationship between specific genes and clone size during viremia, immediate ART, delayed ART, and uninfected conditions. By identifying genes having high clone-size determination scores, we found that T cell exhaustion, T cell activation, and antigen stimulation genes were the underlying transcriptional program that correlated with T cell clone size during delayed ART initiation (as opposed to immediate ART initiation)(Fig. 4e). The upstream regulators of this transcriptional program were LPS, T cell receptor signaling, and cytokine dysregulation (IFNγ, TNF, TGFβ, and IL-21) (Fig. 4f).

**Fig. 4.**
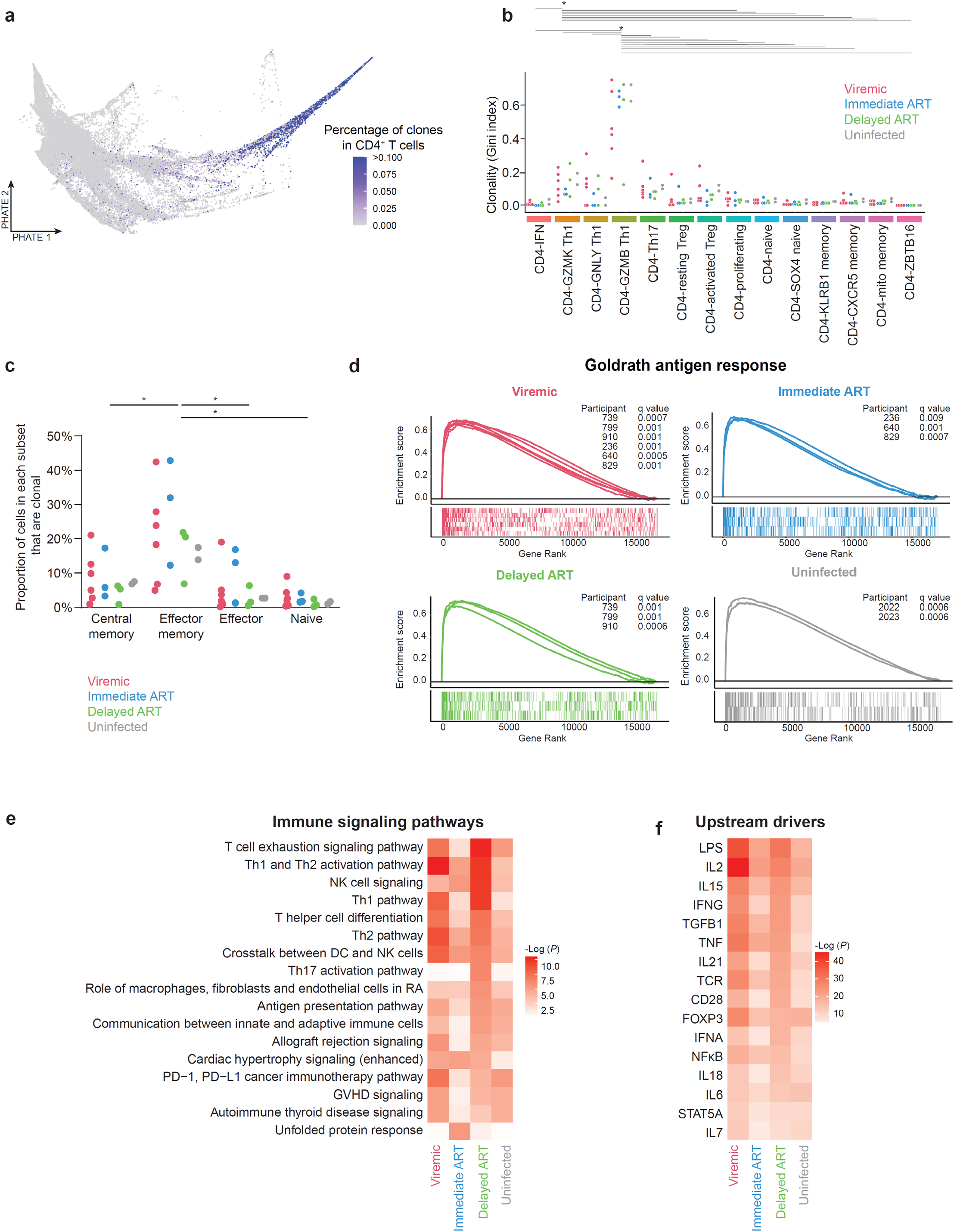
The transcriptional landscape and regulators of T cell clones. **a**, PHATE plot with cells colored by clone size (109,307 cells with TCRs captured) determined by bulk TCR sequencing. **b**, Level of clonality in each transcriptome-defined CD4^+^ T cell clusters measured by Gini Index of cells in that cluster. *P* values were determined by two-way ANOVA followed by Tukey’s multiple comparisons test. *, *P* <0.05. **c**, Proportion of T cell clones in different memory cell subsets. *P* values were determined by two-way ANOVA followed by Tukey’s multiple comparisons test. *, *P* <0.05. **d**, Pearson correlation of gene expression levels and T cell clone size (Log2) determined the rank of genes having expression levels correlating with T cell clone size. **e**, Using elastic net regression, we identified genes that correlated with clone size during viremia, immediate ART, delayed ART, and uninfected conditions. Using these genes for IPA, we identified immune pathways (**e**) and upstream regulators (**f**) that determined T cell clone size. *P* values in e and f were determined by Fisher’s exact test.

### T cell clones harboring HIV-1 RNA^+^ cells remain stable over time

We examined how HIV-1 infection and ART initiation affects T cell clones harboring HIV-1 RNA^+^ cells (clone_HIV-1 HIV-1– HIV-1_^+^) versus T cell clones that do not harbor HIV-1 RNA^+^ cells (clone_HIV-1HIV-1–_). We examined clone_HIV-1_^+^ in addition to HIV-1 RNA^+^ cells (Fig. 3) because T cell clones having the same TCR sequence may proliferate upon the same antigen stimulation, and the immune drivers of the proliferation of these clones may also induce the proliferation of HIV-1 RNA^+^ cells within them. Of note, each clone_HIV-1_^+^ (27 clones, with bulk sizes of 2-1225)only had one or two HIV-1 RNA^+^ cells, either because HIV-1 infection happened after T cell clonal expansion or because other HIV-1-infected cells did not express high levels of HIV-1 RNA without *ex vivo* stimulation. By using shared TCR sequences to track T cell clones, we examined T cell clone size (Fig. 5a–b), T cell expansion dynamics (Fig. 5c– d), and the immune phenotypes of clone_HIV-1_^+^ (Fig. 5e–m), during viremia and after viral suppression. We found that clone_HIV-1_^+^ were significantly larger in clone size than clone_HIV-1 HIV-1–_, during viremia and after viral suppression (Fig. 5a). Immediate versus delayed ART did not significantly change the size of clone_HIV-1_^+^ (Fig. 5b). Despite immediate ART, the size of clone_HIV-1_ ^+^ was larger than clone _HIV-1 HIV-1–_, suggesting that clone _HIV-1_ ^+^ had preferential proliferation despite immediate ART.

**Fig. 5.**
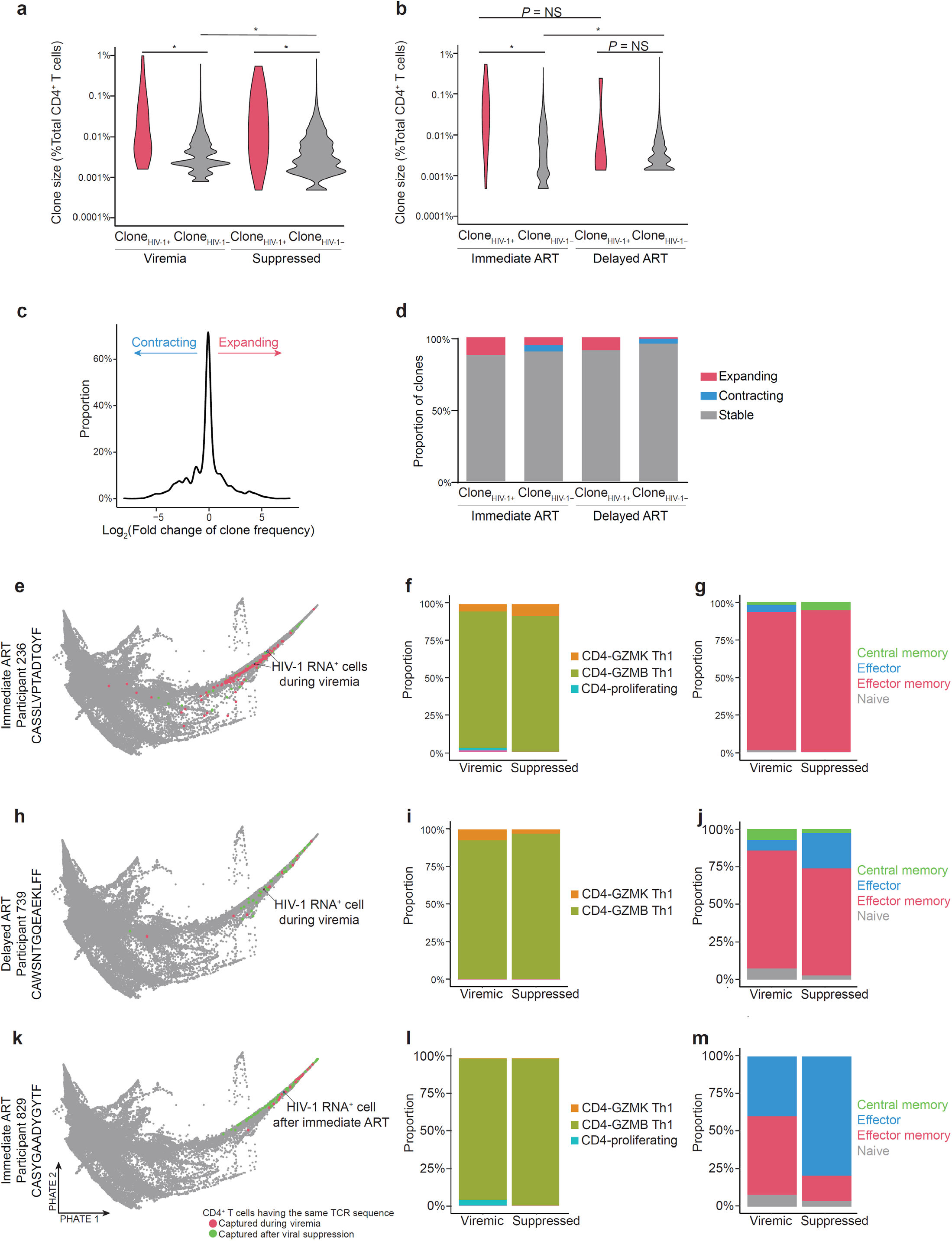
The transcriptional landscape and clonal expansion dynamics of T cell clones harboring HIV-1 RNA_^+^_ cells. **a–b**, Clone size of T cells harboring HIV-1 RNA^+^ cells (clone_HIV-1_^+^, n = 27, 20, and 7 for clones from viremia and suppression, immediate ART, and delayed ART respectively) versus T cells that do not harbor HIV-1 RNA^+^ cells (clone_HIV-1_^−^, n = 12,421, 5,658, and 6,763 for clones from viremia and suppression, immediate ART, and delayed ART respectively respectively). *P* values were determined by Wilcoxon rank-sum test followed by Benjamini-Hochberg procedure. *, *P* <0.05. **c**, Log_2_ fold change of normalized clone size between the viremic and virally suppressed samples. **d**, T cell expansion dynamics in clone_HIV-1_^+^ versus clone_HIV-1_^−^. Significant expansion and contraction was determined by having *P* <0.05 calculated by Fisher’s exact test **e, h, k**, PHATE plots of three representative T cell clones harboring HIV-1 RNA_^+^_ cells (n = 204, 54, and 108 for e, h, and k respectively). **f, i, l**, Transcriptome-defined cluster of T cell clones harboring HIV-1 RNA_^+^_ cells during viremia and after viral suppression in three representative clones. **g, j, m**, Memory phenotype of clone_HIV-1_^+^ during viremia and after viral suppression in three representative clones.

We next examined the change in T cell clone size during viremia and after viral suppression (Fig. 5c). We found that the majority of clone_HIV-1_^+^ remained stable in size and none contracted over time (Fig. 5d), unlike clone_HIV-1–_ which may contract over time (Fig. 5d). We then examined the immune phenotype of clones harboring HIV-1 RNA^+^ cells (Fig. 5e – 5m). We found that the immune phenotype of clone_HIV-1_^+^, as characterized by cluster, did not significantly change over time (Fig. 5f, 5i, 5l), showing few differential expressed T cell function-related genes (Supplementary Fig. 14). These clones were mainly GZMB Th1 cells in these three represented clones. These cells are mainly effector memory cells, and their memory phenotypes did not significantly change over time (Fig. 5g, 5j, 5m).

### Cytotoxic T cell response, antigen response, and cytokine response, shapes the transcription landscape of clone_HIV-1_^+^

We wanted to determine whether clone_HIV-1_^+^ (13 clones with 2 or more cells captured, 4–201 cells in each clone, out of a total of 537 cells, including 276 during viremia and 261 during viral suppression) had a distinct transcriptional signature compare with cells not in clone_HIV-1_ (a total of 78,199 cells, including 45,706 during viremia and 32,493 during viral suppression)(Fig. 6a). To compare the transcriptome profiles of the disproportionate numbers of clone_HIV-1_ (537 cells) with non-clone_HIV-1_^+^ (78,199 cells), we employed single-cell identity definition using random forests and recursive feature elimination (scRFE)^43^ with bootstrapping to identify key genes that were necessary and sufficient to differentiate clone_HIV-1_^+^ from non-clone_HIV-1_^+^. We identified 100 genes (Supplementary Table 8) that were as effective as the 5,000 most highly variable genes for differentiating clone_HIV-1_^+^ from non-clone_HIV-1_^+^ (Fig. 6b, 6c). The number 100 was selected to optimize for the maximum recall without reducing precision in differentiating clone_HIV-1_ from non-clone_HIV-1_^+^ (Supplementary Fig. 15). The majority of the 100 genes identified across all cells were shared between viremic and suppressed conditions suggesting a persistent transcriptional phenotype differentiates clones_HIV-1_^+^ from non-clone_HIV-1_^+^ (Fig. 6d). By examining the differential gene expression profiles of these 100 key genes, we found that the genes that are differentially expressed in clones_HIV-1_^+^ versus clones_HIV-1_^-^ were involved in cytotoxic T cell function (*GZMA, GZMB, GZMH, GNLY, NKG7, CCL4, CCL5, ZNF683, ZEB2, CTSW, CST7*, and *FGFBP2*)^76^ and antigen responses (*HLA-DRB1, HLA-DPA1, HLA-DPB1*, and *FCRL6*) and had lower levels of central memory T cell genes (*TCF7, SELL* (CD62L), *IL7R*, and *LEF1*). To ensure that we were differentiating differences between T cell clones with versus without HIV-1, as opposed to cluster differences such as GZMB Th1 versus non-GZMB Th1, we identified genes that could distinguish clone_HIV-1_^+^ versus cluster-proportion-matched non-clone_HIV-1_^+^ pools (Supplementary Table 8). Among them, 39 genes were significantly upregulated in clone_HIV-1_ (Fig. 6e, Supplementary Table 9). Using gene ontology analysis, we found that these genes were involved in IFNγ responses, cytokine responses, T cell activation, antigen presentation, and viral processes (Fig. 6f – 6p). Among them, we identified cytotoxic CD4^+^ T cell genes (*ZEB2*^77^, *CCL4, ZNF683*)(Fig. 6g – 6i) and antigen response genes (Fig. 6l – 6p). Surprisingly, we identified upregulation of genes that may promote HIV-1 infection, such as *LGALS1* (Galectin-1)(promoting HIV-1 attachment^78^) and *EZR* (Ezrin)(promoting HIV-1 infection^79^). Overall, we found that clone_HIV-1_ demonstrated gene signatures of antigen responses, cytotoxic T cell responses, and cytokine signaling pathways.

**Fig. 6.**
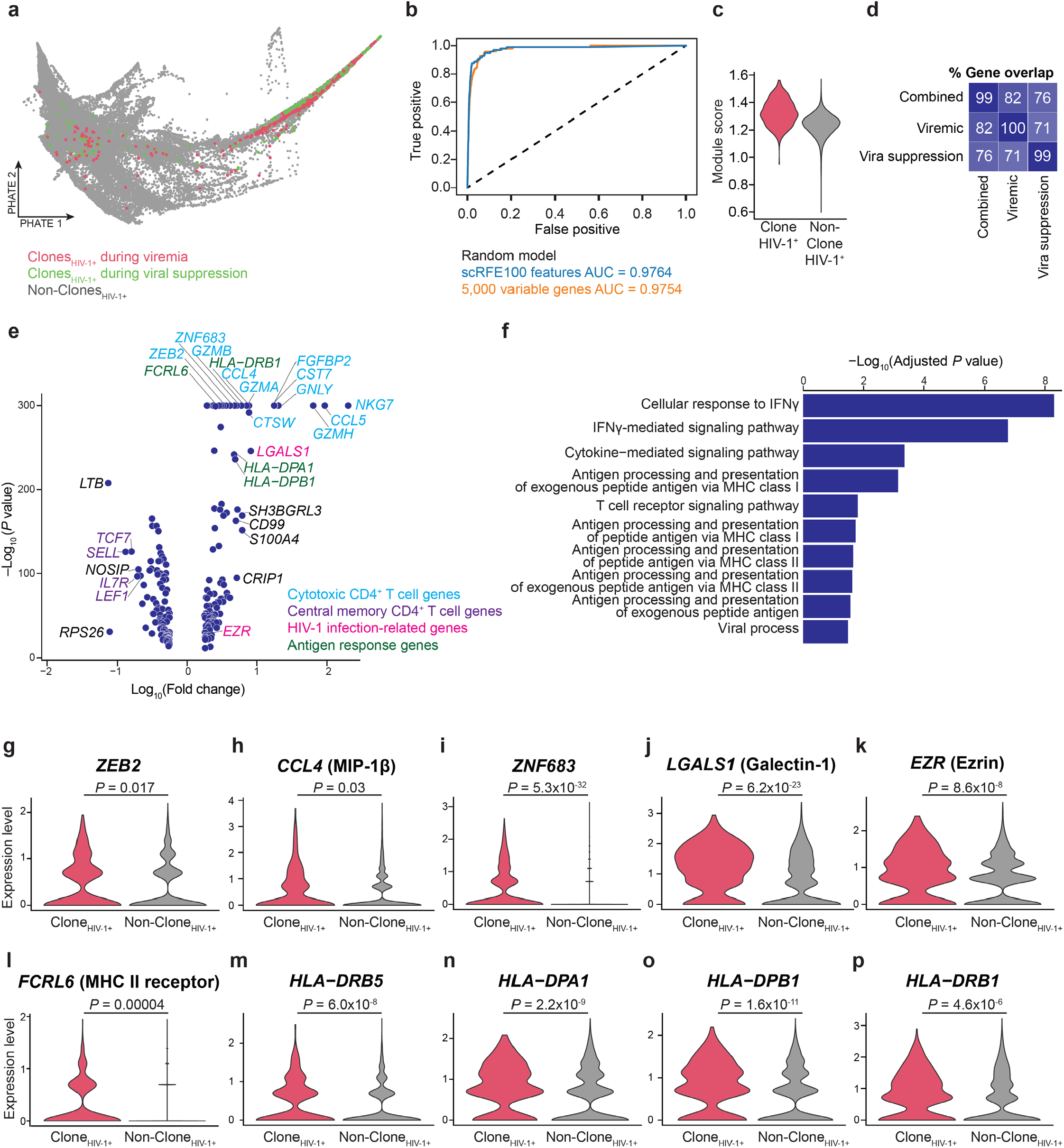
Cellular markers of T cell clones harboring HIV-1 RNA^+^ cells (clone_HIV-1_^+^)(13 clones, 537 cells) versus non-clone_HIV-1_^+^ (78,199 cells). **a**, Cells in clones (≥2 cells) harboring at least one HIV-1 RNA^+^ cell. **b**, Binary classification random forest model performance. Single-cell identity definition using random forests and recursive feature elimination (scRFE) identified key genes that were necessary and sufficient to distinguish clone_HIV-1_^+^ versus non-clone_HIV-1_^+^. We used bootstrapping (comparing the single-cell transcriptome of clone_HIV-1_^+^ to that of random sampling of non-clone_HIV-1_^+^ with 10,000 iterations) to overcome the disproportionate rarity of clone_HIV-1_^+^. We used the top 100 key genes from these iterations to train a model and tested how confident that these 100 genes, as opposed to top 5,000 highly variable genes, or random chance, can differentiate cells in clone_HIV-1_^+^ versus non-clone_HIV-1_^+^. AUC, area under curve. **c**, Module score of the 100 key genes in clone_HIV-1_^+^ versus non-clone_HIV-1_^+^. **d**, Gene overlap of the 100 key genes identified by scRFE bootstrapping analysis when trained on all cells, cells from the viremic time point, and cells from the suppressed time point. **e**, Volcano plot of differential gene expression analysis of the 100 key genes comparing clone_HIV-1_^+^ to non-clone_HIV-1_^+^ using differential expression testing in Seurat (Wilcoxon rank-sum test). **f**, Gene ontology enrichment of genes (n = 39) from the 100 key genes that are significantly upregulated in clone_HIV-1_^+^ relative to a cluster proportion matched pool of non-clone_HIV-1_^+^ using Enrichr. **g–p** Gene expression differences of key genes upregulated in clones containing an HIV-1 RNA_^+^_ cell relative to a cluster proportion matched pool of clone_HIV-1_^−^ (n = 573 and 3,305 for clone_HIV-1_^+^ and non-clone_HIV-1_^+^, respectively). Y axis shows natural log expression levels.

## Discussion

We found immune drivers that promote HIV-1 persistence and cellular markers as potential therapeutic targets. First, we found that an ongoing TNF response was the major immune dysfunction in delayed versus immediate ART (Supplementary Fig. 7–8), shaped the transcriptional program of HIV-1 RNA^+^ cells (Fig. 3h), and was an upstream regulator shaping T cell clonal expansion (Fig. 4f). Second, we found that cytotoxic CD4^+^ T lymphocytes, particularly those expressing *GZMK* (granzyme K) and *GZMB* (granzyme B), harbored HIV-1 RNA^+^ cells (Fig. 3b) and clone_HIV-1_^+^ (Fig. 4). Third, we found that T cell clones harboring HIV-1 RNA^+^ cells were larger in clone size (Fig. 5a). Fourth, using machine learning algorithms, we identified 22 markers for HIV-1 RNA^+^ cells (Fig. 3) and 39 upregulated genes for clone_HIV-1_^+^ that distinguished HIV-1 RNA^+^ cells from HIV-1 RNA^−^ cells and clone ^+^ from non-clone_HIV-1_^+^. Altogether, we found that HIV-1 resides in cytotoxic T cells which are naturally proliferative. These cytotoxic CD4^+^ T cells continue to be impacted by antigen stimulation and TNF responses from viremia through viral suppression.

Our study highlights that immune dysfunction is established rapidly during acute HIV-1 infection. Unless initiated in hyperacute infection^34^, early ART initiation could not fully revert chronic immune activation. By identifying co-regulated genes during HIV-1 infection, we found that acute viremia induced IFN, antigen, and TNF responses (Fig. 2, Supplementary Fig. 7–9). Both immediate and delayed ART reduced IFN responses to the level in uninfected individuals, but antigen responses and TNF responses persisted (Supplementary Fig. 7). Importantly, reducing TNF responses was the most prominent impact that immediate ART provides relative to delayed ART, although immediate ART did not fully restore TNF response to the level in uninfected individuals (Supplementary Fig. 7 and 8), particularly on GZMK Th1 cells (Fig. 1k, 2j)(see below). Immediate versus delayed ART did not significantly change the cell subsets harboring HIV-1 RNA^+^ cells (Fig. 3) and the clonal expansion dynamics of T cell clones in CD4^+^ T cells (Fig. 4) and in clone_HIV-1_^+^ (Fig. 5). Pathway analysis suggested that both antigen responses and TNF signaling shaped the landscape of HIV-1 RNA^+^ cells (Fig. 3h) and T cell clonal expansion (Fig. 4e). Of note, despite the strong effect of antigen responses on T cell clonal expansion (Fig. 4d), common γ chain cytokines that promote homeostatic proliferation (such as IL-2 and IL-15) shaped the transcriptional landscape of HIV-1 RNA^+^ cells (Fig. 3h) and T cell clones (Fig. 4f), suggesting that both antigen stimulation and homeostatic proliferation^12^ contribute to HIV-1 persistence. Strategies that can reduce chronic antigen stimulation in people with HIV-1^80,81^, or strategies targeting HIV-1-induced cytokine dysregulation^62^, should be explored to eliminate these immune drivers that promote the persistence of HIV-1-infected cells.

Antigen-responding CD4^+^ T cells are known to be an important source of the HIV-1 latent reservoir^9-11^. Antigen-responding CD4^+^ T cells have distinct functional states, such as cytolytic granule secretion (such as granzyme B, granzyme K, and perforin), cytokine production (such as IL-2, IFNγ, and TNF), and antiviral chemokine production (such as CCL3, CCL4, and XCL1). Cytotoxic CD4^+^ T cells are critical effectors responding to HIV-1^82-86^, cytomegalovirus^87,88^, SARS-CoV-2, and tuberculosis^89^ infections. We have previously found that HIV-1 RNA^+^ cells after *ex vivo* stimulation exhibit Th1 polarization and upregulate antiviral chemokines *CCL3, CCL4, and XCL1*^*13*^. Our study identified an under-appreciated role of *GZMB* and *GZMK*-expressing cytotoxic CD4^+^ T cells as a source of HIV-1 reservoir. First, we found that GZMK Th1 CD4^+^ T cells harbored HIV-1 RNA^+^ cells (Fig. 3b). The GZMK Th1 cells were phenotypically distinct from GZMB Th1 cells (Fig. 1). Importantly, *GZMK*-expressing CD4^+^ T cells were recently found to be important in SARS-CoV-2 infection^59^ and in anti-tumor cytotoxicity^50^. Second, we found that GZMB and GZMK Th1 T cells exhibited strong antigen responses (Fig. 1h), were clonally expanded (Fig. 4b, Supplementary Fig. 5f), were enriched of HIV-1 RNA^+^ cells (Fig. 3b), and were the major T cell phenotype of clone_HIV-1_^+^.(Fig. 5). Third, cytotoxic CD4^+^ T cell markers were among the key cellular markers that could distinguish HIV-1 RNA^+^ cells from HIV-1 RNA^−^ cells and clone_HIV-1_^+^ from non-clone_HIV-1_^+^, such as *GZMK, CST7*, and *CCL4* (Fig. 3i, 3k, 6e). We postulate that despite ART, persistent antigen stimulation and TNF responses induce the proliferation of HIV-1-infected cytotoxic CD4^+^ T cells and promote HIV-1 persistence.

Specific targeting of HIV-1-infected cells requires viral or cellular markers that can distinguish HIV-1-infected cells from uninfected cells. Studying HIV-1-infected cells from people with HIV-1 provides the most clinically relevant understanding^13,19,21,23,80,90-92^. We identified HIV-1 RNA^+^ cells and clone_HIV-1_^+^ without *ex vivo* stimulation and identified genes that marked these cells. Using machine-learning analysis, we found that HIV-1 RNA^+^ cells and clone_HIV-1_^+^ can be distinguished by a distinct transcriptional phenotype. While the majority of identified markers were consistent with T cell activation during viremia, a subset of markers were previously unreported, such as *GZMK, CLIC1, NEAT1*, and *C9orf16* for HIV-1 RNA^+^ cells (Fig. 3) and *ZEB2, ZNF683*, and *LGALS1* for clone_HIV-1_^+^ (Fig. 6). The combination of markers may define useful populations for studying HIV-1-infected cells in the future. These cells, though not latent at time of capture, are a source of persistent immune activation through HIV-1 transcriptional activity. Our finding raises the possibility of targeting HIV-1 RNA^+^ cells and clone_HIV-1_^+^ more precisely.

## Acknowledgements

We thank all study participants. We thank Alex K. Shalek, Sam Kazer, Joseph Craft, Steven Kleinstein, Nicolas Chomont, and Michael Betts for their insightful suggestions. This work is supported by Yale Top Scholar, Rudolf J. Anderson Fellowship, NIH R01 AI141009, NIH R61 DA047037, NIH R37 AI147868, NIH R01 DA051906, NIH UM1 DA051410, NIH CHEETAH P50 AI150464, NIH BEAT-HIV Delaney Collaboratory UM1 AI126620, Gilead HIV-1 Research Scholar Grant, American Foundation for AIDS Research (amfAR 110029-67-RGRL), Lupus Research Alliance – Celgene, and NIH T32 AI055403 (J.A.C.). The Sabes and MERLIN studies were supported by NIH/NIDA R01 DA032106 and NIH/NIDA R01 DA040532 (A.D.). We gratefully acknowledge ART drug donation from Merck & Co. and Gilead Sciences Inc.

## Author contributions

J.A.C. and Y.-C.H. designed the experiments, performed analyses, and wrote the manuscript. J.A.C. performed experiments and bioinformatic analyses. J.A.C. performed machine learning analyses with help from N.P. and D.V.D. D.P.-S., S.P., C.G., R.A., J.L., J.C., S.S., and A.C.D. recruited study participants and processed clinical samples. A.D. designed the Sabes and MERLIN studies and assumed oversight for study conduct and analysis.

